# Curiosity, latent learning, and cognitive maps

**DOI:** 10.1101/2020.05.31.123380

**Authors:** Maya Zhe Wang, Benjamin Y. Hayden

**Affiliations:** Department of Neuroscience, Center for Magnetic Resonance Research, and Center for Neuroengineering, University of Minnesota, Minneapolis MN 55455

**Author notes:** Corresponding author: Maya Zhe Wang, Department of Neuroscience and Center for Magnetic Resonance Research University of Minnesota, Minneapolis MN 55455.

## Abstract

Curiosity refers to a desire for information that is not driven by immediate strategic or instrumental concerns. Latent earning refers to a form of learning that is not directly driven by standard reinforcement learning processes. We propose that curiosity serves the purpose of motivating latent learning. Thus, while latent learning is often treated as an incidental or passive process, in practice it most often reflects a strong evolved pressure to consume large amounts of information. That large volume of information in turn allows curious decision makers to generate sophisticated representations of the structure of their environment, known as cognitive maps. Cognitive maps facilitate adaptive and flexible behavior while maintaining its adaptivity and flexibility via map updates based on new information. Here we describe data supporting the idea that orbitofrontal cortex (OFC) and dorsal anterior cingulate cortex (dACC) play complementary roles in curiosity-driven learning. Specifically, we propose that (1) OFC tracks the innate value of information and incorporates new information into a detailed cognitive map; and (2) dACC tracks the environmental demands and information availability to then use the cognitive map for guiding behavior.

## Introduction

The natural environment offers a plethora of rewards to most foragers but acquiring these rewards requires knowledge – for example, trees with ripe apples may be inhomogeneously distributed in a forest (Calhoun and Hayden, 2015; Gottlieb and Oudeyer, 2018; Mobbs et al., 2018; Oudeyer and Smith, 2016). Consequently, natural decision-makers are faced with a constant overwhelming poverty of information. Any steps they can take to alleviate that poverty, even fractionally, offers great survival benefits relative to rivals. This information gap is an area where psychological and neuroscientific tasks used in the laboratory differ from those that occur in the natural environment and that presumably drove our evolution. In the laboratory, research subjects (both human and non-human) are typically given all information except that which is experimentally relevant. As a consequence, there are areas where failure to consider the paucity of valuable in our evolved environment can mislead us.

One of those areas is curiosity. Curiosity, which we can define as a drive for non-strategic information, has long fascinated psychologists (Berlyne, 1966; Oudeyer et al., 2016; Kidd and Hayden, 2015; Byrne, 2013; Loewenstein, 1994; Golman and Loewenstein, 2018). It is a major driver of learning and determinative of the success of development; it applies to humans and to non-human animals (ibid.). Recent work has begun to demonstrate its preserved features across species and over the lifespan. It appears to be associated with at least somewhat discrete neural circuits (Gruber et al., 2014; Lau et al., 2020; Cervera et al., 2019). While there is no universally accepted definition of the term, curiosity, a common feature of most definitions is that it involves a demand for information that is intrinsic non-strategic, and/or non-instrumental. This definition distinguishes curiosity from information seeking in, for example, the well-understood bandit task (Cohen et al., 2007). In that case, information-seeking, and subsequent learning, is driven by rewards.

### Latent learning

Classical concepts of learning held that all learning is driven by reinforcement contingencies. Formally, the presence or lack of reward or punishment promotes the positive or negative associations between actions and the results they produce, or between different stimuli. These ideas are fundamental to the “Law of Effect” and are fundamental to the idea of Hebbian learning (Thorndike, 1927; Walton et al., 2010). That work, in elaborated form, is central to reinforcement learning, one of most successful psychological theories and the basis of a generation of systems neuroscience. Notably, these concepts in turn form the basis of much of modern machine learning.

However, even as early as 1929, it was shown by Hugh C. Blodgett, a graduate student of Edward Tolman at UC Berkeley, that robust learning could occur in the absence of reward (Blodgett, 1929). This makes sense to anyone who can still effortlessly recite the lyrics to advertising jingles from their childhood. But it was the source of much debate in psychology departments in the first half of the 20th century. Later members of Tolman’s lab followed up on this work, which is summarized in Tolman’s classic paper (Tolman, 1948). In this paper, Tolman characterized the simpler reinforcement learning approach as the “*stimulus response school*,” and described the idea of cognitive maps, which he believed depended on latent learning.

In a classic latent learning setup, a rat is released into a large maze with no reward. Naive rats typically amble around the maze, ostensibly with no purpose. According to behaviorist theories of the stimulus-response school, the rat should not learn anything in particular about the layout of the maze because no reward or punisher was provided. Later, in a different experimental session, the experimenters can introduce a reward to a specific location in the maze. The same rats, on encountering the reward, would then be given a chance to run the maze again from the beginning. The rats with maze exposure learned to locate the reward much more quickly than ones who were naive to that maze. Tolman coined the term “latent learning” to describe the learning of maze layout by rats when they were not reward motivated. To drive this point home, a later experiment provided food and water in the maze and rats went through latent learning when they are satiated (not motivated). They showed high efficiency using the learned maze layout to find food or water when later they visited the maze hungry or thirsty (Tolman, 1948; Spence et al., 1950). Tolman proposed that these rats had learned (latently) the layout of the maze without any stimulus-reward association, a theory that violated then-dominant dogmatic versions of behaviorism. Specifically, Tolman proposed that latent learning allowed the rats to form a cognitive map of the layout of the maze.

### Curiosity and cognitive maps

Blodgett’s rats did not move randomly throughout the maze for no reason. Their decision to move was not costless, nor was the mental effort involved in observing and consolidating those observations into a mental map. Any forager placed within a complex natural environment must naturally trade off between the costs and benefits of exploration. In addition to the metabolic costs of locomotion, sensory processing, and learning, active exploration carries opportunity costs: that time could be better spent searching for food, courting and reproducing, or avoiding predators. The decision-maker, subject inevitably to extremely high selection pressures, does not wander idly. But we also can use any standard extrinsically rewarded economic explanation to motivate latent learning. Even motivational processes driven by distal reward seeking must necessarily discount future rewards and uncertain rewards, and the benefits of exploration are unavoidably delayed beyond the temporal horizon and, individually, infinitesimally unlikely. So reward-maximizing calculation is unlikely to motivate search. Evolution must step in and endow the decision-maker with intrinsic motivation to learn (Bennett et al., 2016; Grant et al., 1998; Tversky and Edwards, 1966; Kidd and Hayden, 2015).

Curiosity is likely to be especially important for learning cognitive maps. Cognitive maps refer to detailed mental representations of the relationship between various elements in the world and their sequelae. Having a cognitive map allow a decision-maker to not just guess what will happen but also to deal with unexpected changes in our environment. The classic idea about cognitive maps - also attributable to Tolman - is that they allow us to respond flexibly when the layout of a maze was later changed (Tolman, 1948; Tolman et al., 1946). That kind of flexibility - the ability to have insight - is very difficult to implement with basic reinforcement learning processes (Schoenbaum and Roesch, 2005; Wilson et al., 2014). Instead, we must have a sophisticated representation of the structure of the world, and the ability to navigate counterfactuals and unexpected changes.

Cognitive maps require a level of detail that is not normally available from reinforcement learning processes. Specifically, cognitive maps involve a detailed representation of the linkages between adjacent spaces, and allow for vicarious travel along those linkages. That representation, then, is a level of abstraction higher than the simple stimulus-outcome-action mapping that standard reinforcement learning gives. But because it is so much richer, and more detailed, it requires orders of magnitude more information than standard reward-motivated reinforcement learning can give. Getting that information cannot occur if it needs extrinsic rewards - those rewards simply are not available in the environment. It requires intrinsic rewards.

We propose that what Tolman observed as latent learning, was motivated and enabled by curiosity and information-seeking behavior (free, non-motivated exploration in his case). However, Tolman conceived of latent learning as a fundamentally passive process, one that took place during apparently purposeless exploration - almost as if by accident. We propose, instead, that latent learning in practice tends to be more actively driven. However, this purposive exploration must be e driven by the evolutionary advantage brought by curiosity and ultimately by the extreme information gap experienced by foragers in the natural world.

### The analogy to artificial intelligence

The problems faced by a naturalistic decision-maker or forager are similar in many ways to the problems faced by artificially intelligent (AI) agents in machine learning problems. Consider for example that AI can be trained to perform a classic Atari games using straightforward RL principles (Mnih et al., 2013). In these cases, the agent must learn a strategy using an elaborated gradient descent procedure – not different from how the stimulus-response school imagined rats learned a maze. But those games, especially the ones that AI is good at, like Video Pinball and Breakout, differ from natural situations in key ways that make them easy for RL algorithms. In contrast, the real world – and some games like Pitfall and Montezuma’s Revenge - are what is known as hard-exploration problems (Bellemare et al., 2016; Ecoffet et al., 2018; Ecoffet et al., 2019). In such problems, rewards are sparse (they require dozens or hundreds of correct actions), so gradient descent procedures are nearly useless. For example, in Pitfall, the first opportunity to gain any points comes after ∼60 seconds of perfect play involving dozens of precisely timed moves. Moreover, rewards are often deceptive (they result in highly suboptimal local minima, so getting a small reward promotes adherence to a suboptimal strategy). Finally, real-world problems often present very abstract goals (e.g. “design an attractive furniture layout for the study on a medium budget”). RL agents that do well at relatively naturalistic hard-exploration games tend to have deliberate hard-coded exploration bonuses.

The AI domain provides a good illustration of how cognitive maps can be crucial for the success of curiosity. The optimal search strategy in sparse (natural) environments is typically to identify a locally promising region and then perform strategic explorations from that spot to identify subsequent ones (Ecoffet et al., 2020). That exploration will not be random, but will take place along identified high-value destinations. AI agents suffer from the problem of *detachment*, that is, when they explore the environment, they leave the relatively high-reward areas of space to explore lower-reward ones. Most such areas are likely to be dead ends, and, when a dead end is detected, the agent ought to return to the high reward area and pursue other promising paths. However, the basic curiosity-based approach, which gives intrinsic rewards for novelty, repel the agent from returning to the promising region of space, precisely because it’s the most familiar and least intrinsically rewarded (it’s also not extrinsically rewarding, because any extrinsic reward has been consumed on the path there, and does not replenish in the meantime). This in turn requires making some kind of internal map of space so that the agent can return to the locus of high potential reward and explore more efficiently than a wholly random path. One example would be a bee hunting for promising hive sites. That bee must be able to have some map of the local environment (Cheeseman et al., 2014).

A closely related problem that AI agents - and real-world agents as well - face, is the problem of *derailment*. To explore a space efficiently, an agent must be able to return to promising states and use those as a starting point for efficient exploration. From there, the agent must engage in random search. However, in real environments, returning to a promising state may require a very precise sequence of actions that cannot be deviated from - so stochasticity must be controlled until that state is achieved, at which case it must begin again in earnest. As such, stochastic search must be carefully controlled depending on one’s place in the larger environment - which requires basic mapping functions, and cannot be done with simple RL-type learning. Moreover, important factors governing the exploration process, such as detecting an information gap, deriving the value of information itself, and directing exploration towards potential sites that might be low in external reward but high in information/entropy, simply cannot be supported by only experienced reward history. The key to achieve this is to have a mental map, or internal model, of what is available, and what is novel and potentially offer high information content (high entropy).

An intuitive example is a monkey learning to harvest fruit at the top of a tree. The tree is tall and with lush leaves such that the fruits are not visible from the safety on the ground. While monkeys need the calories and nutrients to survive and thrive, and the fruits offer both, the climb is treacherous and the cost of falling is high. So in order to maximize reward and minimize cost, but also being able to make better and better predictions on which new branches to go to for potentially more fruits and/or a better view to detect them, the monkey must have some kind of map of its internal state in order to strategically deploy search.

### Operational definition of curiosity

Developing these ideas about curiosity, latent learning, and cognitive maps holds great potential in neuroscience. However, it faces several problems from the get-go. We and others have defined curiosity as a motivation to seek information that lacks instrumental or strategic benefit (Kidd and Hayden, 2015; Loewenstein, 1994; Berlyne, 1966; Oudeyer et al., 2016). By this definition, many explorative and playing behaviors qualify as a demonstration of curiosity (Glickman and Sroges, 1966; Byrne, 2013). From the perspective of scientific research, this definition is problematic because it is vague and so does not readily lend itself to the laboratory.

In an effort to remedy these drawbacks, we developed an operational definition that combines three criteria: (1) a curious research subject is willing to sacrifice primary reward in order to obtain information; (2) the amount of reward a subject is willing to pay scales with the amount of potentially available additional information; and (3) additionally gained information provides no obvious instrumental or strategic benefit. This definition is deliberately conservative; that is, it excludes much of behavior that is likely to be curiosity-driven.

This definition justifies the observing task as a laboratory instrument for manipulating curiosity, although that task is subject to certain criticisms that have not yet been resolved (e.g. Beierholm and Dayan, 2010). On the other hand, formalized risky choice tasks offer a straightforward opportunity to understand preferences with close control of information (Heilbronner, 2017; Heilbronner and Hayden, 2016). We therefore devised a more complex task that would circumvent published criticisms of the observing task (Wang and Hayden, 2019). This task is based on the observation that monkeys seek counterfactual information - information about what would have happened had they chosen differently. In the *counterfactual curiosity task*, monkeys choose between two risky offers. Behavior is overtrained (hundreds of thousands of trials), reducing the chance that monkeys have erroneous theories about payoff distributions. During testing, monkeys are sometimes given the opportunity to choose an option that will provide valid information about the outcome that would have occurred had they chosen the other option. Monkeys are willing to pay to choose this option, indicating that they are curious about counterfactual outcomes. Moreover, monkeys paid more for options that provide more counterfactual information. We speculate that this curiosity-driven information-seeking helps monkeys to develop a sophisticated cognitive map of their task environments.

### Functional neuroanatomy of curiosity in the frontal lobes

Our ultimate goal is to understand the neural circuitry underlying curiosity-driven choice. Here we summarize the tentative picture, with a focus on two prefrontal regions, the orbitofrontal cortex (OFC) and the dorsal anterior cingulate cortex (dACC). Both regions are implicated in neuroimaging studies of curiosity (Charpentier et al., 2018, Kobayashi and Hsu, 2019; Van Lieshout et al., 2018). The neuroanatomy of curiosity is more complex and includes other areas such as hippocampal areas (Kang et al., 2009; Gruber et al., 2014; Jepma et al., 2012) and basal ganglia (White et al., 2019). But we would like to highlight OFC and dACC for their potential involvement that bridges curiosity, latent learning, and cognitive maps. Briefly, we propose that the OFC and dACC have complementary roles and that these roles are best described as acquisition and update of the cognitive maps and instrumental application of cognitive maps, respectively.

#### Orbitofrontal cortex

We propose that OFC serves to track the innate value of information, to maintain a cognitive map of state space, and to update that map when new information is gained. The clear role of OFC in cognitive mappings has been one of the major intellectual advances of the past decade, and is demonstrated in rodent, monkey, and human species. (Wilson et al., 2014; Wang and Hayden 2017; Wikenheiser and Schoenbaum, 2016; Schuck and Niv, 2016). OFC also carries various contextual variables not directly related to economics (Roesch et al., 2006; Feierstein et al., 2006; Wallis et al., 2001; Sleezer et al., 2016). Our lab’s contribution to this comes from a study using a variant of the observing task (Blanchard et al., 2015). We found that OFC neurons encoded the value of information and (confirming much previous work) the value of offers. Critically, however, OFC used distinct codes for informational value and for more standard juice value - highlighting the special role of information – it was not an additive value to the primary reward, but of its own distinct category. this distinction is possibly for completing and updating the cognitive map of the task/environmental structure -- even though this information may not offer strategic benefit in any near future. In other words, OFC doesn’t use a single coherent value code across contexts, but rather, represents task-relevant information in multiple formats, as would be expected in a map rather than a simple reinforcement learning situation. Of course, OFC does not achieve this alone. Studies using similar paradigms discovered that information is signaled by other systems, including the midbrain dopamine system (Bromberg-Martin and Hikosaka, 2009 and 2011;Guru et al., 2020).

### ACC: regulation of information flow in the cognitive map

We propose that dACC plays a distinct and complementary role of OFC. Specifically, it appears to track both information delivery and level and task demands for use by OFC in updating the cognitive map and applying it to instrumental use. This idea is motivated by the observation that dACC tracks informativeness, counterfactual information (White et al., 2019, Hayden et al., 2009), environmental demands (Kolling et al., 2012; Hayden et al., 2011), as well as various economic variables (e.g. Bush et al., 2002; Seo and Lee, 2007; Azab and Hayden, 2017, 2018). It is further motivated by observations about the relative hierarchical positions of the two regions and the relative contributions to choice (Rushworth et al., 2011; Hunt et al., 2018; Yoo and Hayden, 2018).

In a recent study using a Pavlovian paradigm, White et., al. (2019) trained monkeys to associate juice rewards with various reward probabilities with different fractals. Then, information (or non-information) stimuli followed the fractals. Single units in dACC showed increased firing rates to increased uncertainty, and thus to higher expectation of information (when the uncertainty resolved). Moreover, dACC firing rates ramped up to the anticipation of the information that came with the resolution of the uncertainty. In other words, dACC neurons did not simply encode different levels of uncertainty which remained at a constant level for each trial; nor did they ramp up firing rates in anticipation to reward delivery These results showed that dACC tracks the lack of information and anticipates the delivery of information. Our own results paint a similar picture (Wang and Hayden, 2020). Using the observing task, we find that, following uninformative choices, firing is enhanced until information delivery and is scaled with the amount of information gained through resolving gamble uncertainty. Moreover, in our study, enhanced dACC firing also correlated with higher probability of information-seeking choices later in the trial.

### Conclusion and future directions

Curiosity is a strong motivator of learning, development, and choice. It has long been treated mystically, as if it is impenetrable to scholarly study. This attitude is found, for example, in some of the research that treats curiosity as a human-specific phenomenon. Even when treated as a regular psychological phenomenon, curiosity is often studied in an ad hoc manner. That approach was necessary for early studies, but more recent work has made great progress in developing formal approaches to understand the phenomenon systemically study its neural substrates. That formal approach, aided by remarkable progress in AI, has in turn allowed neuroscientists to tentatively start to understand the circuity of curiosity. That work in turn will likely be critical for understanding naturalistic decision-making, which is marked by the need to make quick decisions with orders of magnitude less information than would be idea.

## METHODS

### General Methods

All animal procedures were performed at the University of Rochester (Rochester, NY, USA) and were approved by the University of Rochester Animal Care and Use Committee. All experiments were conducted in compliance with the Public Health Service’s Guide for the Care and Use of Animals. Two male rhesus macaques (*Macaca mulatta*), aged 9-10 years and weighting 8.0-9.9 kg served as subjects. Both subjects had extensive previous experience in risky decision-making tasks. Subjects had full access to food (LabDiet 5045, ad libitum) while in their home cages. Subjects received at minimum 20 mL per kg of water per day, although in practice they received close to double this amount in the lab as a result of our experiments. No subjects were sacrificed or harmed in the course of these experiments.

Visual stimuli were colored rectangles on a computer monitor (see Figure 1). Stimuli were controlled by Matlab with Psychtoolbox. Eye positions were measured with Eyelink Toolbox. A solenoid valve controlled the delivery duration of fluid rewards. Eye positions were sampled at 1,000 Hz by an infrared eye-monitoring camera system (SR Research, Osgoode, ON, Canada). A small mount was used to facilitate maintenance of head position during performance.

**Figure 1.**
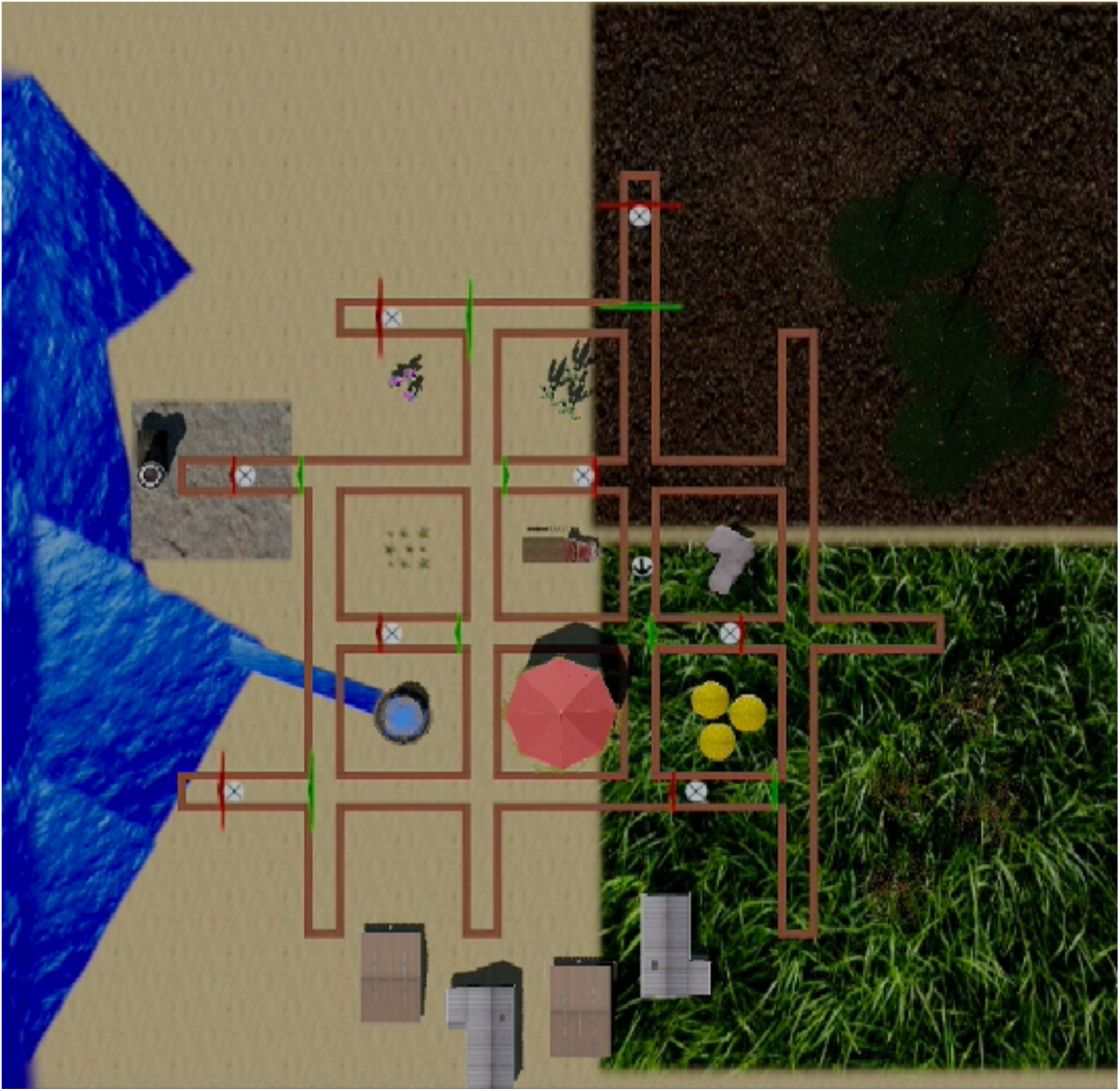
A virtual maze our lab has used for monkeys based on the classic 14-unit T-alley maze of Tolman (Elliott, 1928). The difference here is that the current veirtual maze cannot be expanded into a linear track in contrast to the original maze. Tolman and his graduate students placed rats in mazes like this one and found that they would explore the maze unrewarded and would demonstrably learn the features of the structure of the maze in the absence of rewards, a result that is difficult to explain using then-dominant simple stimulus-response learning theories. Tolman proposed that the rats generated a cognitive map that instantiated features of the maze and could be consulted to drive flexible behavior.

**Figure 2.**
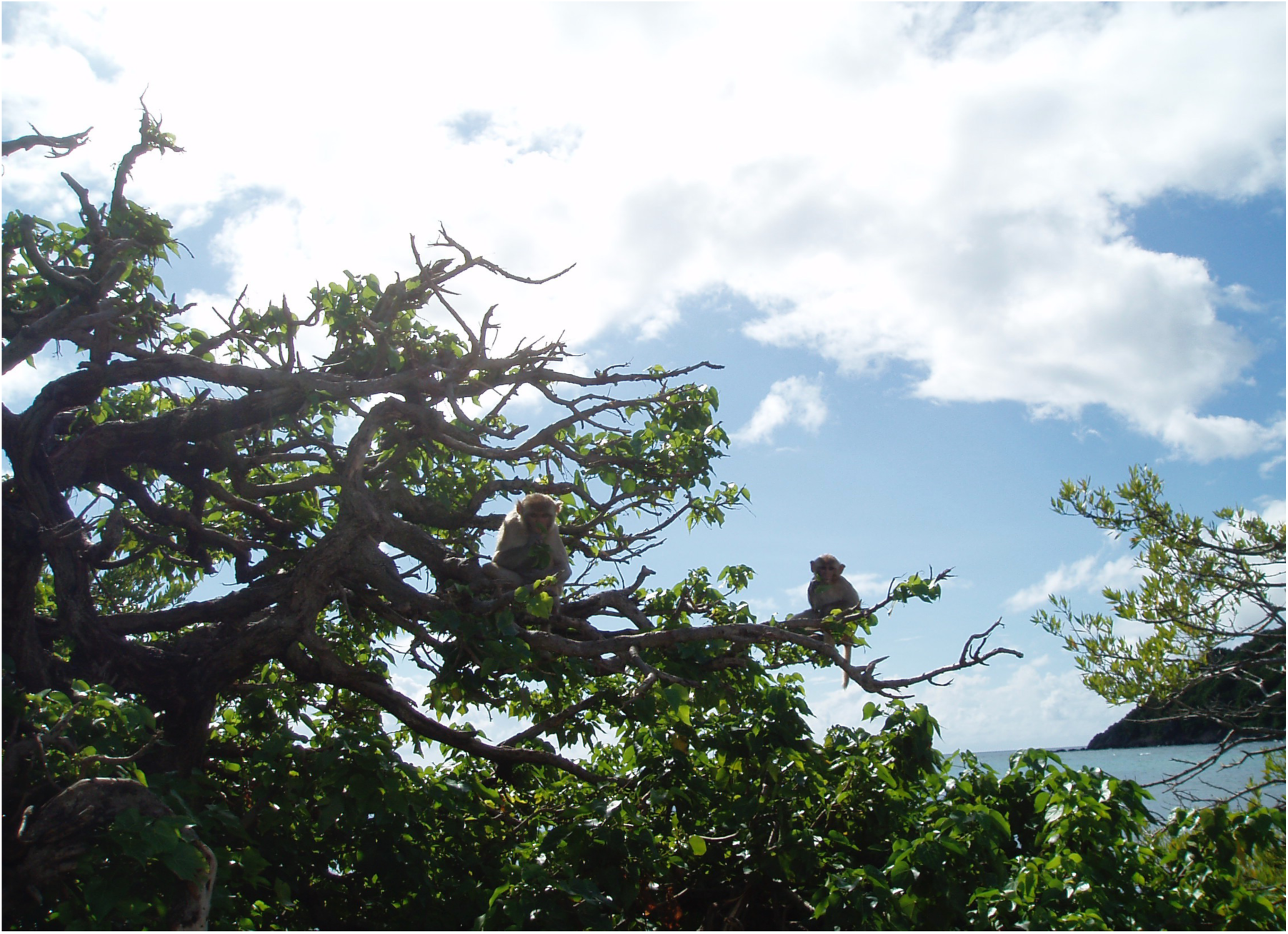
Monkey in a tree, illustrating the problem of *derailment* in curiosity research. The monkey must learn foraging strategy through trial and error, which requires highly variable exploration of the environment. But getting to the end of a branch is somewhat risky and requires suppressing stochastic variability. To successfully deploy curiosity the monkey must have a cognitive map of where variability is good and where it is bad.

**Figure 3.**
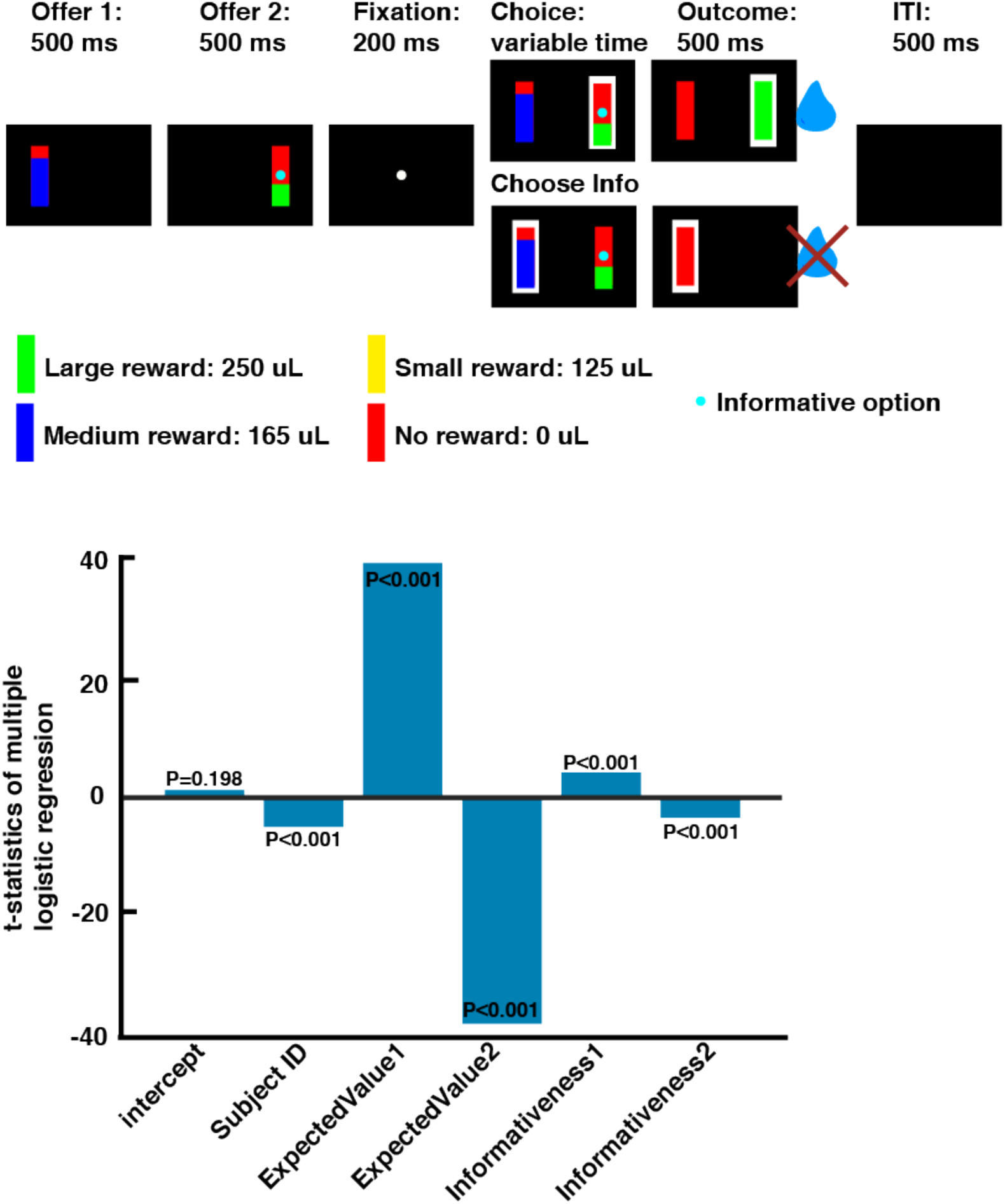
In our curiosity task, subjects could choose between risky options for juice rewards. In some trials, they could also gain information about what would have occurred had they chosen differently. By analyzing preference curves on such trials, we could quantify their subjective value of counterfactual information. We found a small but significant positive valuation of counterfactual information in both subjects tested.

**Figure 4.**
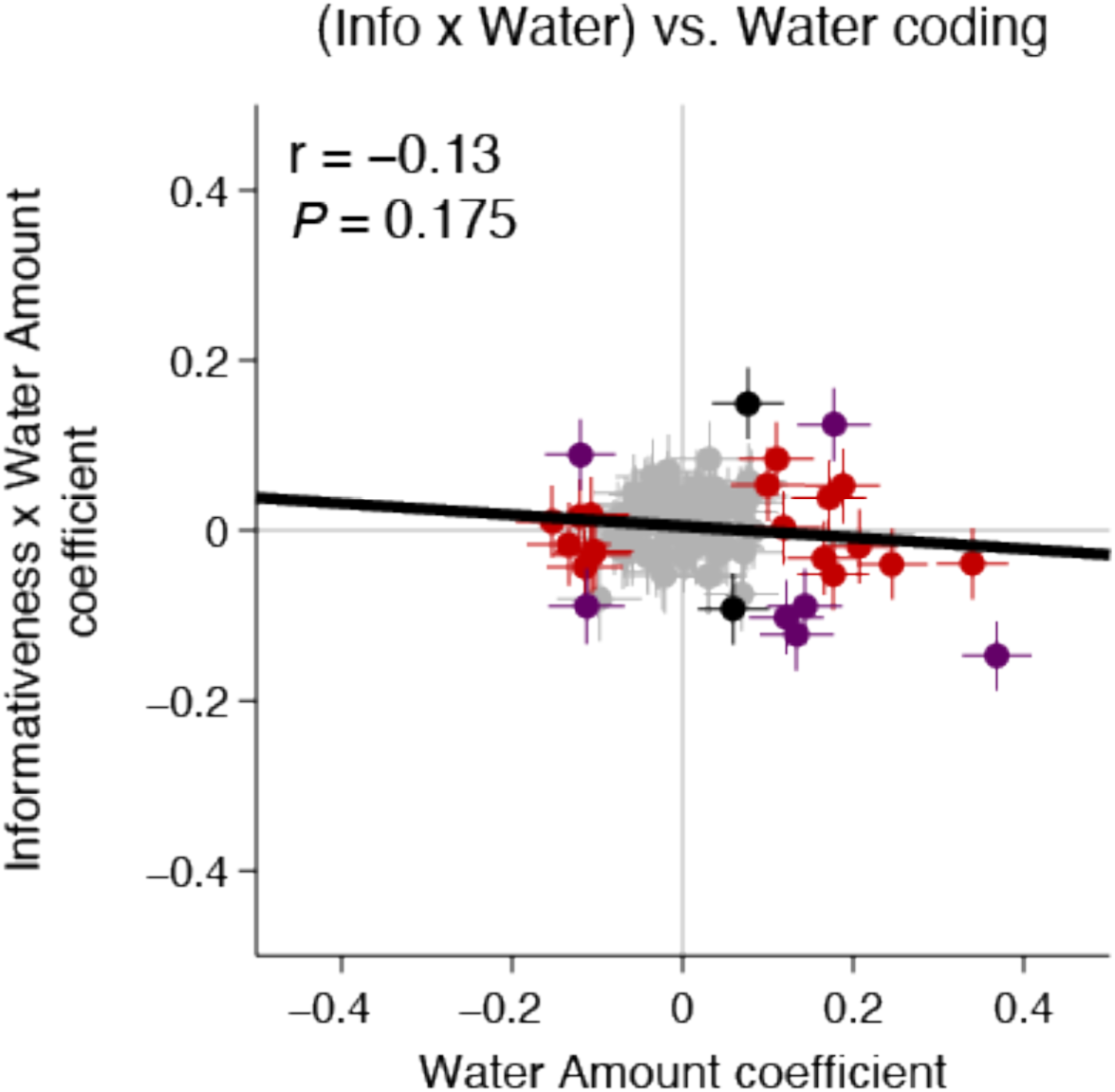
In our observing task, responses of an ensemble of OFC neurons to offers varying in informational value and reward value. We find that individual neurons encode both variables (horizontal and vertical axes indicate tuning coefficients for the two dimensions respectively). However, those codes are themselves uncorrelated, as indicated by the lack of a significant slope between the two dimensions.

Subjects had never previously been exposed to decision-making tasks in which counterfactual information was available.

### The Counterfactual Information Task

The task structure was a close variant of a general one that we have used many times in the past. Subjects fixated, in sequence, on two options, located on the two sides of the computer monitor. They had been extensively trained (5+ years in both cases) on tasks like this in the past and were adept at making effective choices in those.

Two subjects (B and J) performed a novel task designed to measure preference for counterfactual information (Fig. 1a). Due to the extensive exposure to similar tasks and the simplicity of the current task, no pre-training was used. Both subjects were trained directly on the current task and achieved above 80% accuracy within the first three days of training. Following completion of additional training, we collected 8142 trials of behavior from both subjects (5086 trials from subject B and 3056 trials from subject J). On each trial, subjects chose between two randomly selected gambles presented asynchronously on the left and the right side of the screen. Gambles were represented by rectangular visual stimuli and differed in three dimensions: payoff, probability, and informativeness. Payoff came in three sizes, small (125 microliters), medium (165 microliters), and large (250 microliters), each corresponding to a yellow, blue, and green portion of the rectangle, respectively. Probabilities were randomly drawn from a uniform distribution between 0 and 1 (101 steps; step size 0.01). The height of the yellow/blue/green portion of the rectangle indicated the probability of winning the gamble and the height of the red portion indicated the probability of losing (that is receiving no reward for that trial). Informativeness of a gamble was indicated by a cyan dot on the center of the rectangle for an informative option and the lack of a cyan dot for a non-informative one. The informative option promised valid information about the payoff that would have occurred had the alternative option been chosen. Probability, payoff, and informativeness were independently randomized on each trial. On 50% of the trials, only one option was informative (*info choice* trials). On 25% of the trials, both options were informative (*forced info* trials). The forced info trials were equivalent to what would be called full-feedback trials in human judgment and decision-making literature (Camilleri & Newell, 2011). On the remaining 25%, neither option was informative (*no info* trials). These are equivalent to what are called partial-feedback trials.

We have previously used this general structure (without the informativeness manipulation) to probe macaques’ preferences for uncertainty. Critically, via controls, we have demonstrated that macaques treat these stimuli as if they provide explicit information about the structures of gambles.

Each trial started with the appearance of offer 1 (500 ms) followed by a blank 500 ms delay. Offer 1 position was randomized for each trial. Then offer 2 appeared on the other side of the screen (500 ms) followed by another 500 ms delay. After a 200 ms fixation, both gambles appeared on the screen and subjects chose the preferred option by shifting gaze to it and maintaining that gaze for 200 msec. Subsequently, if an informative option was chosen, gamble outcomes for both offers were resolved. If a non-informative option was chosen, the gamble outcome for only the chosen offer was resolved. Resolution of a gamble involved filling the gamble rectangle with the payoff color while delivering a water reward (if the gamble result was win), or filling the gamble rectangle with red color and delivering no reward, (if the gamble result was a loss). The outcome epoch lasted for 800 ms and was followed by a 1000 ms inter-trial interval (ITI) and then the start of next trial.

Consider, for example, a subject performing the following trial. First, offer 1 appears on the left side of the computer monitor. It is a non-informative option (it has no cyan dot) with 80% probability (indicated by the height of the blue section) of yielding a medium reward (165 uL, indicated by blue color) and 20% probability of yielding no reward (indicated by red color). After a second, offer 2 appears. Offer 2 is an informative option (it has a cyan dot) that corresponds to a 45% probability (indicated by height of green segment) of yielding 250 uL (indicated by green color), and 55% probability of getting no reward (indicated by red color). After a brief pre-choice eye fixation, the subject chooses offer 2. This choice resulted in a win with water reward delivery and the presentation of counterfactual outcome information.

## Acknowledgements

We thank Ethan Bromberg-Martin for helpful discussions.

